# Activity-dependent lateral inhibition enables the synchronization of olfactory bulb projection neurons

**DOI:** 10.1101/2024.06.24.600470

**Authors:** Tal Dalal, Rafi Haddad

## Abstract

Information in the brain is represented by the activity of neuronal ensembles. These ensembles are adaptive and dynamic, formed and truncated based on the animal’s experience. One mechanism by which spatially distributed neurons form an ensemble is by synchronizing their spike times in response to a sensory event. In the olfactory bulb, odor stimulation evokes rhythmic gamma activity in spatially distributed mitral and tufted cells (MTCs). This rhythmic activity is thought to enhance the relay of odor information to the downstream olfactory targets. However, how specifically the odor-activated MTCs are synchronized is unknown. Here, we demonstrate that optogenetic activation of one set of MTCs can gamma-entrain the spiking activity of another set. This lateral synchronization was particularly effective when the recorded MTC fired at the gamma rhythm, facilitating the synchronization of only the substantially active MTCs. Furthermore, we show that lateral synchronization did not depend on the distance between the MTCs and is mediated by granule-cell layer neurons. In contrast, lateral inhibition between MTCs that reduced their firing rates was spatially restricted to adjacent MTCs and was not mediated by granule-cell layer neurons. This dissociation between these two interaction types suggests that they are mediated by different neural circuits. Our findings propose a simple yet robust mechanism by which spatially distributed neurons entrain each other spiking activity to form an ensemble.

**Highlights:**

1. MTC activation entrains the spike timing of other MTCs in an activity-dependent and distance-independent manner.
2. MTC to MTC suppression is activity- and distance-dependent
3. Spatially distributed Granule cell layer neurons control MTC’s spike timing, yet do not substantially affect their odor-evoked firing rate.

## Introduction

Information in the brain is represented by the activity of ensembles of neurons, typically interconnected through a network of interneurons (Buzsáki and Chrobak, 1995). These interactions reshape the ensemble activity as it evolves in time and space. One mechanism by which spatially distributed neurons form an ensemble is by synchronizing their spiking activity in response to a sensory event (Buzsáki, 2010). This synchronization is thought to enhance the transmission of information to downstream targets (Dalal and Haddad, 2022; Macleod et al., 1998; Pritchett et al., 2015; Sohal, 2016).

Olfactory processing starts with the activity of odorant-activated olfactory sensory neurons. The axons of these sensory neurons terminate in one or two anatomical structures called glomeruli located on the surface of the olfactory bulb (OB). Each glomerulus is innervated by several MTCs, which then project the odor information to several cortical regions. Different odors activate different olfactory receptors. The odor-activated MTCs interact with each other via a dense network of inhibitory neurons spanning all OB layers (Burton, 2017). Several studies have shown that odorants evoke strong spike gamma-entrainment in MTCs (Beshel et al., 2007; Dalal and Haddad, 2022; Fukunaga et al., 2014; Lepousez and Lledo, 2013), as well as synchronous firing of MTCs (Doucette et al., 2011; Kashiwadani et al., 1999). This synchronization is likely to be mediated by granule cells (GCs) (Schoppa, 2006) located in the granule cells layer (GCL) and has recently been shown to enhance odor-information transmission to the Piriform cortex (Dalal and Haddad, 2022). However, the mechanism by which specifically the odor-activated MTCs are synchronized is unknown. One prominent hypothesis is that GCs enable synchronization solely between odor-activated MTCs via an activity-dependent mechanism for GABA-release (Egger and Kuner, 2021; Lage-Rupprecht et al., 2020).

GCs are the most abundant type of neuron in the OB. They form dendrodendritic synapses with MTCs (Shepherd, 2004) and, via these synapses, can provide recurrent and long-range lateral inhibition. Earlier studies that were not aware of the role of EPL interneurons in regulating MTC spiking activity, suggested that GCs mediate lateral inhibition between MTCs that can suppress their firing rates (Arevian et al., 2008; Giridhar et al., 2011; Yokoi et al., 1995). In contrast, more recent studies demonstrated that in-vivo lateral suppression of MTC firing rate is sparse and weak (Fantana et al., 2008; Lehmann et al., 2016; Pressler and Strowbridge, 2017). Consistent with this weak effect on MTC firing rates, one study showed that optogenetically silencing the GCs did not affect odor-evoked inhibitory responses (Fukunaga et al., 2014). These findings cast doubt on the role of GCs in suppressing MTCs’ odor-evoked firing rate (Burton, 2017).

Here, we used odor and optogenetic stimulations of MTCs and GCL neurons in anesthetized mice, to study how active MTCs interact to regulate their spikes timing and firing rates. We found that MTCs form two dissociated types of interactions. One interaction enables the synchronization of only activated MTCs dispersed on the OB surface and is mediated by GCL neurons. The other type of interaction affecting MTCs firing rate is limited to relatively nearby MTCs, and, contrary to some common views in the field, is not mediated by GCL neurons.

## Results

### Activity-Dependent Lateral Synchronization of MTCs

To investigate how MTCs interact we expressed the light-gated channel rhodopsin (ChR2) exclusively in MTCs by crossing the Tbet-Cre and Ai32 mouse lines (Grobman et al., 2018; Haddad et al., 2013), and extracellularly recorded the spiking activity of MTCs in anesthetized mice during optogenetic stimulation using tungsten electrodes. We first mapped each recorded cell’s receptive field, i.e., the set of MTCs on the dorsal OB that affect its firing rates when light-stimulated. Pseudo-random light patterns composed of multiple light spots were projected over the OB surface (Supplementary Figure 1a) and the spike-triggered-average was computed to obtain the receptive field map (Figure 1a; ‘STA’, see Methods). Light patterns comprised of 5-10 multiple light spots of size 88-110 µm^2^. This stimulation protocol ensured the activation of varied MTC combinations associated with different glomeruli, among them the MTC we recorded from. As expected, light-stimulating the area above the recording electrode activated the recorded MTC (the ‘hotspot’). This protocol also revealed regions on the OB surface that had a suppressive effect on the activity of the recorded MTC (Figure 1a). Based on the receptive field map, we selected several spots and light-stimulated them either alone or simultaneously with the ‘hotspot’ using four increasing light intensities (Figure 1a-b; N = 5 mice; 27 well-isolated single units; overall 127 pairs were tested, 2-8 different pairs per cell; see Methods for spots selection criteria). We found that in a subset of neurons, paired light activation precisely aligned the spike times of the recorded MTC across trials (Figure 1c), giving rise to a gamma rhythm (Figure 1d-e). To quantify the change in spikes gamma entrainment at the population level, we computed the difference in the area under the power spectrum density (PSD) curve between the two conditions at the gamma range (40-70Hz, Supplementary Figure 1b; see Methods). We found that the change in entrainment was significantly higher than zero (Figure 1f; P < 0.001 and P = 0.13 for real and shuffled data, respectively; two-tailed paired *t*-test, N = 319/511 values from all pairs and light intensities that significantly responded to light stimulation). Overall, we found that in ∼16% of the stimulated pairs, paired light-activation significantly enhanced the gamma entrainment (N = 50/319 values that exceeded the 95% confidence interval of the shuffled distribution, colored in brown in Figure 1f). Varying the light intensities revealed that the enhanced gamma entrainment was most effective when the postsynaptic MTC firing rate was ∼40Hz (Figure 1g). Analyzing the locations in which paired activation significantly increased the temporal precision revealed no significant correlation between the pair’s distance and its gamma entrainment change, suggesting that lateral entrainment is driven by MTCs across the glomerular map without any discernable spatial organization (Figure 1h; r = 0.05, P = 0.73, Spearman correlation). Overall, we found activity-dependent enhancement of MTC spike precision in the gamma range that is distance-independent.

**Figure 1:**
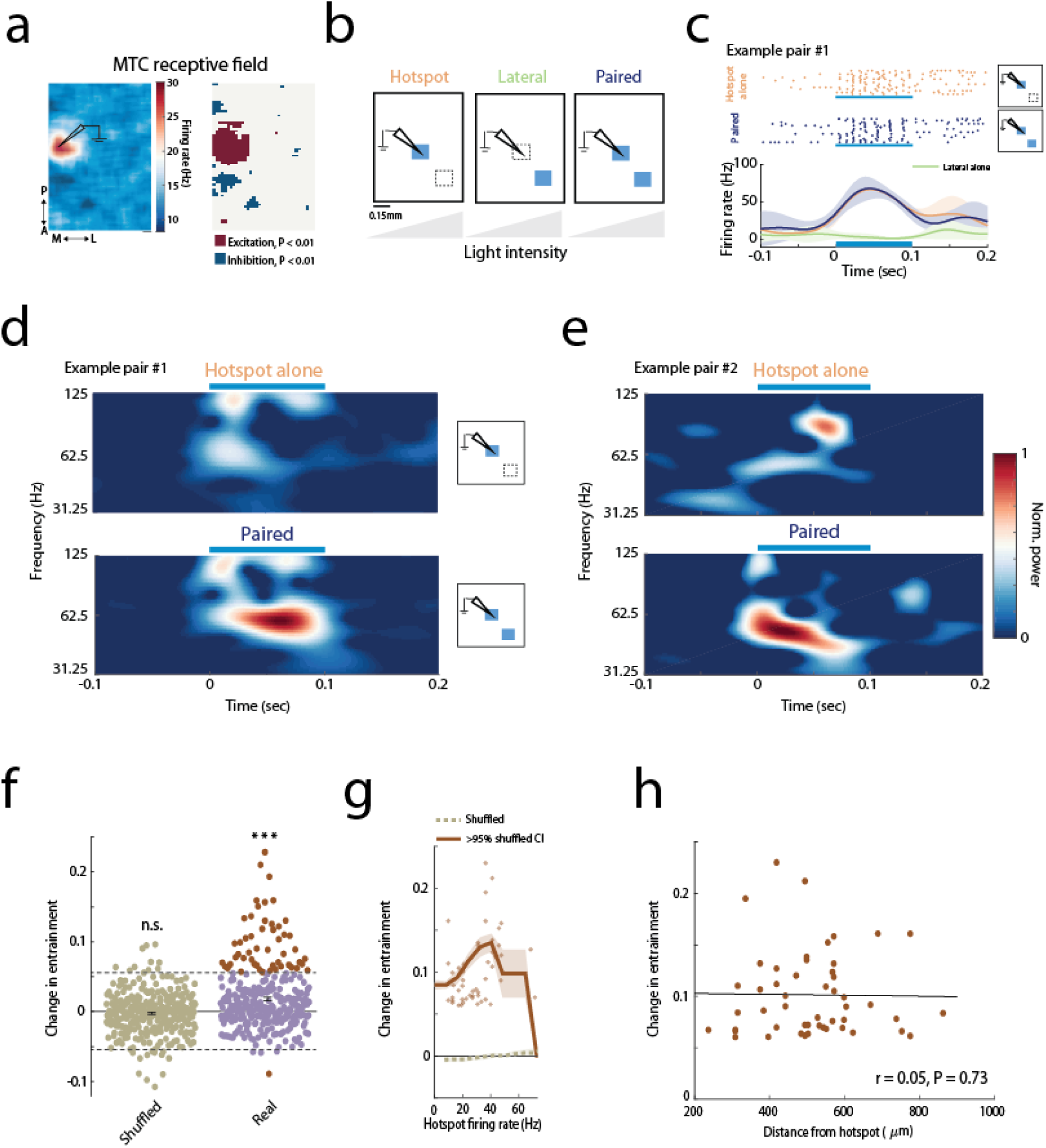
Activity-dependent lateral-entrainment of spike times a) Left: STA map with an excitatory region near the electrode location and presumably several surrounding inhibitory spots. Right: the significance map (P < 0.01 relative to shuffled data, see Methods). Scale bar, 0.1mm. b) Schematic illustration of the experimental setups. Photo-stimulation of each spot alone (hotspot or lateral spot conditions marked by orange and green text, respectively) or paired stimulation (marked in blue) using four different light intensities. c) Raster plots and smoothed PSTHs of the response to light stimulation of the hotspot (top, orange) and paired stimulation (middle, blue). Note, the increase in spike time accuracy within and across trials when both spots are activated (middle panel). Paired light stimulation did not affect the average firing rate in this example (lower panel). The effect of light stimulation of the lateral spot alone is shown in green. d-e) Examples of MTCs time-frequency wavelet analysis from two different mice. Example pair #1 is the pair displayed in **c**. Both examples show a strong gamma rhythm following paired stimulation. In pair #1, gamma power peaked at ∼58Hz, and pair #2 at ∼48Hz. f) Paired stimulation increases spikes’ temporal precision. Mean ± SEM of the change in spikes entrainment at the population level (N = 319/511 values from all pairs and light intensities that significantly responded to light stimulation, P = 0.13 and P < 0.001 for shuffled (green) and real (purple) data, respectively; two-tailed paired *t*-test). In brown are values that exceeded the 95% confidence interval of the shuffled data distribution values of increased and decreased spike entrainment, respectively; confidence interval is marked by dashed black lines). g) Lateral entrainment is activity-dependent. The moving average of the data shown in **f** is plotted as a function of the firing rate of the postsynaptic MTC (N = 50 values of increased entrainment). The increase in entrainment was largest when the neuron fired at ∼40Hz. The color code is the same as in **f**. The shuffled data is shown in a dashed green line. h) Spike entrainment does not depend on the distance between the MTC pair. No significant correlation was found between the increase in spike-entrainment and the distance from the hotspot (r = 0.05, P = 0.73, Spearman correlation; N = 50 values with significant increase in spike entrainment, brown dots in **g**).

**Supplementary Figure 1:**
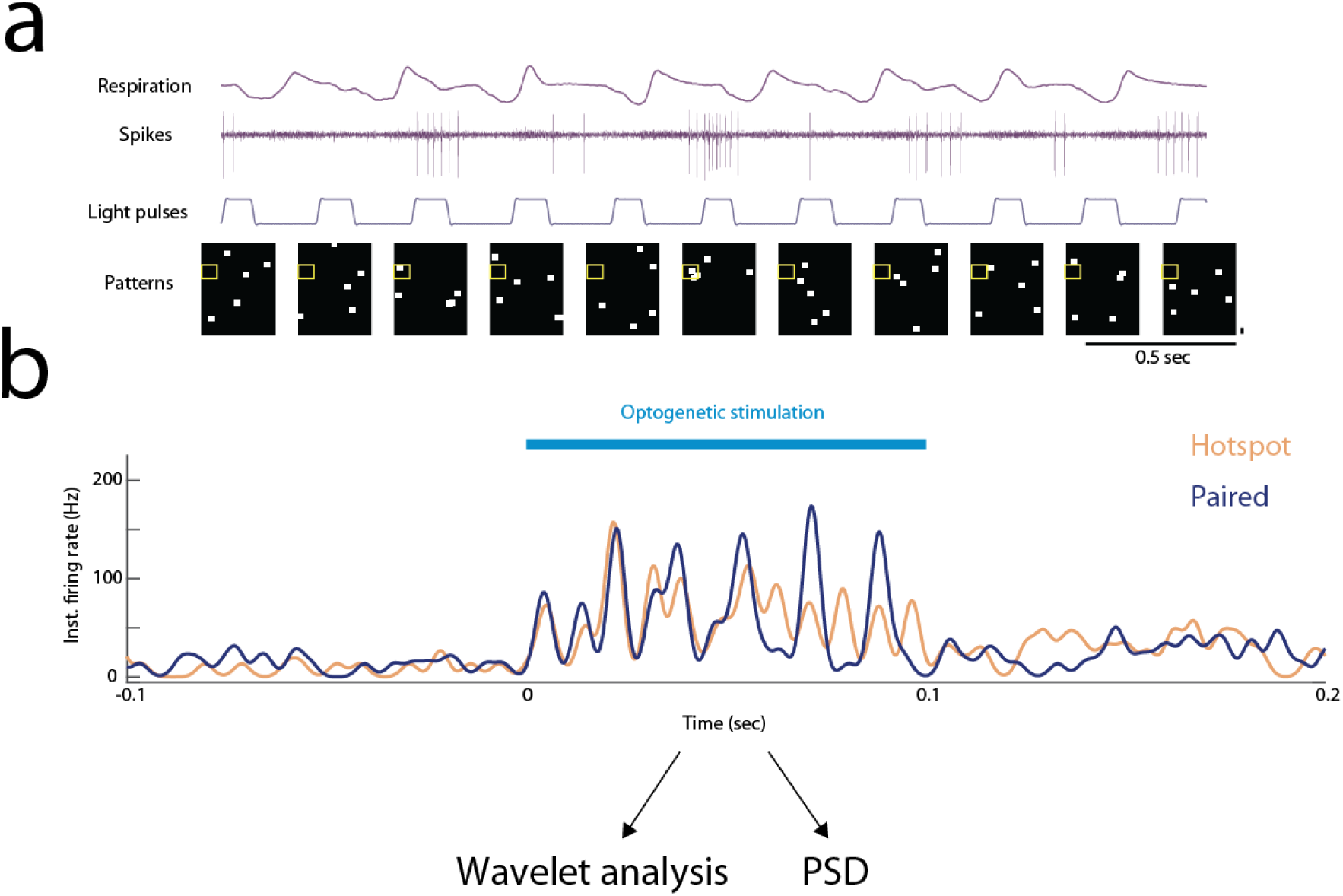
Paired MTC activation induces spikes entrainment a) Light stimulation protocol. An image containing N randomly distributed light patches (N = 5 in this example) is projected on the dorsal bulb in each trial. The light patterns are shown in the bottom panel. The yellow rectangle marks the region around the recording electrode. The spiking activity and the respiratory signal are shown above. Multiplying each pattern by the firing rate it evoked and averaging across all trials gives the STA activity map (see Methods). Scale bars: 0.5 second; 110 µm. b) A description of the analysis used to compute the spikes entrainment. For each condition (hotspot alone or paired activation) a PSTH was computed using a Gaussian window of 2 ms. We then fed this PSTH into the wavelet analysis (as shown in Figure 1d-e, see Methods) and computed the power spectral density (PSD) using a Multitaper analysis during the stimulus presentation (100ms, see Methods).

### Activity-Dependent Lateral Suppression of MTCs is Confined in Space

In addition to lateral synchronization between co-active MTCs, we found that paired activation could suppress the recorded MTC firing rate (termed here as lateral suppression, Figure 2a-b). Figure 2c shows the response of a recorded MTC when light-stimulated alone and simultaneously with other MTCs under four different light intensities (pair #1 or pair #2, as marked in Figure 2a). Activation of pair #1 evoked lateral suppression that was dependent on the recorded MTC firing rate, while activation of pair #2 did not affect the MTC firing rate for the four light intensities tested . Plotting the evoked change in firing rate caused by paired stimulation as a function of the MTC firing rate when stimulated alone revealed that lateral suppression is most effective when the recorded MTC fires in the gamma range (i.e., ∼30-80 Hz, Figure 2d). Light stimulating the lateral spots alone did not affect the MTC baseline firing rate (Supplementary Figure 2a). Overall, in 19% (24/127) of the tested MTC pairs, the recorded MTC firing rate was significantly suppressed during paired stimulation compared to hotspot stimulation alone. In contrast to the lack of spatial organization between MTC pairs that caused spike entrainment, we found a significant correlation between the distance of the paired spots and the level of evoked suppression (Figure 2e; r = 0.45, P = 0.001, Spearman correlation), consistent with a recent study (Peace et al., 2024). To further verify that lateral inhibition is limited to proximal MTCs, we analyzed the inhibition found in the receptive field (STA) maps. Centering all z-scored maps relative to the ‘hotspot’ location (N = 27 neurons from 5 mice) revealed that MTC-to-MTC suppressive interactions are strongest and densest in regions that are adjacent to the recorded MTC (∼400 µm) and diminished beyond that (Figure 2f and Supplementary Figure 2b). Furthermore, computing the STA maps while excluding the light patterns that contained a light spot that hit the electrode location (‘hotspot’ area) resulted in maps that did not contain suppressive regions (Supplementary Figure 2c-d). This analysis further confirms that lateral suppression requires the recorded MTC to be active. In summary, we show that MTC-to-MTC suppressive interactions are spatially confined and activity-dependent.

**Figure 2:**
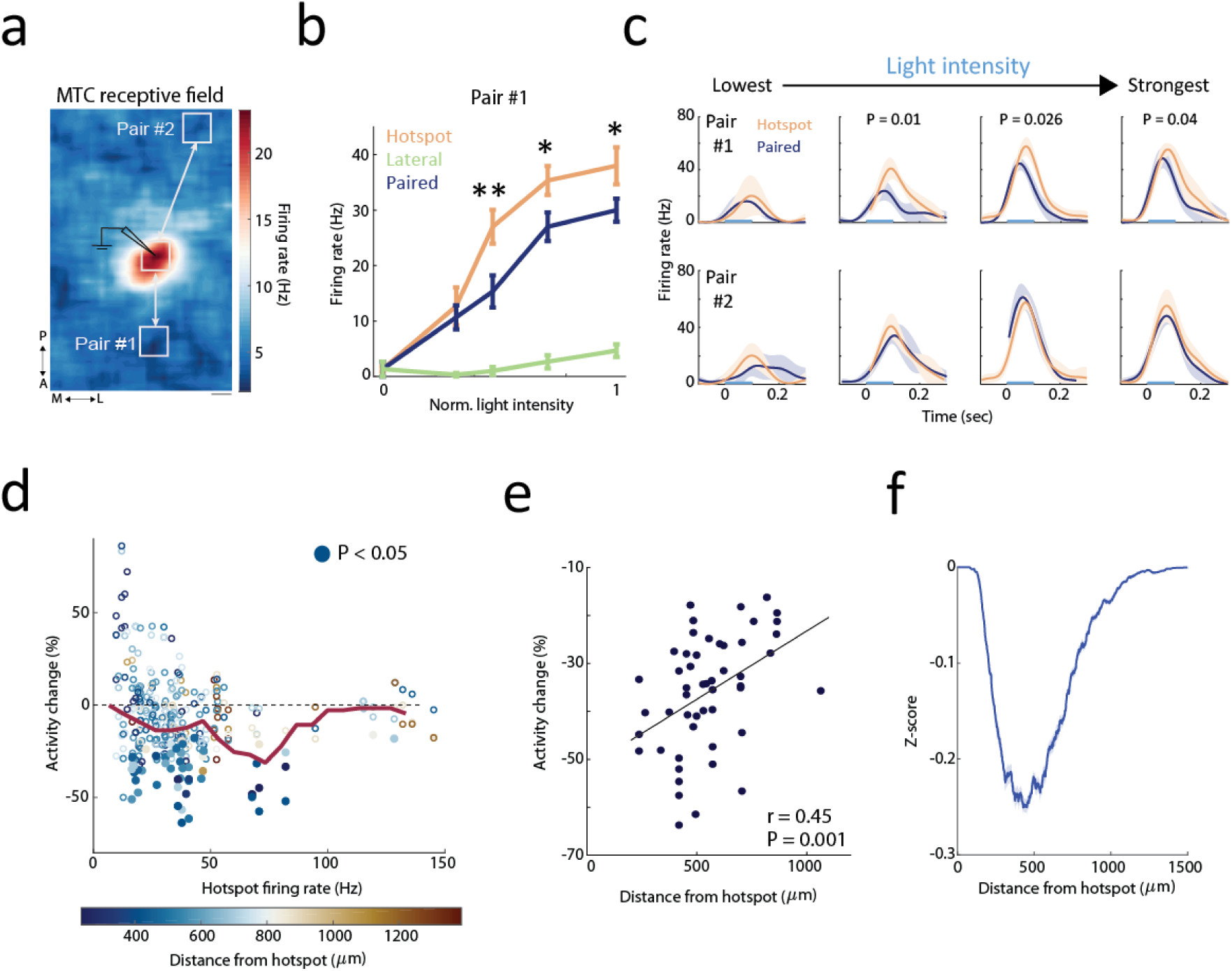
Activity-dependent lateral suppression of MTCs is confined in space a) An example of a MTC receptive field (STA map). The white rectangles mark the spots exposed to light stimulations. The hotspot location is marked with an electrode drawing. Scale bar, 0.1mm. b) Mean ± SEM firing rates of light stimulating pair #1 from **a** for each of the three conditions across all four light intensities. Activation of pair #1 (blue) caused a reduction in the recorded MTC firing rates only when the recorded MTC fired above ∼25 spikes/sec. Zero denotes the baseline firing rate. *P < 0.05, **P < 0.01, two-tailed paired *t*-test. c) PSTHs of pair #1 and #2 responses to light stimulations at four different light intensities. Paired stimulation of pair #1 evoked activity-dependent suppression. Paired stimulation of more distant neurons in pair #2 did not affect the recorded MTC firing rates for all tested light intensities. Light stimulation is marked with a blue bar (0.1 sec). PSTHs without a p-value showed no significant change between the firing rates of paired stimulation and hotspot stimulation alone. d) Summary analysis of the effect of paired activation on MTCs firing rate. Each point marks the percentage change in firing rate (Y-axis) relative to the firing rate elicited by light stimulating the hotspot alone (X-axis). Suppressive effects (i.e., negative activity change) occurred mainly when the MTC fired in the gamma range (∼30-80 Hz). Color code denotes the distance between the light-activated MTC pair. Filled circles mark significant activity change (P < 0.05, two-tailed unpaired *t*-test, N = 51/319 data points). The red line shows the moving average. Only light intensities that elicited a significant light response were analyzed (319/511; P < 0.05, two-tailed paired *t*-test). e) Lateral suppression degrades with distance. Spearman correlation between the change in firing rate for all significant inhibitory pairs (the filled circles in **d,** N = 51) and their distance to the hotspot (r = 0.45, P = 0.001). f) Z-score Mean ± SEM change in MTC activity across all STA maps superimposed and centered relative to the ‘hotspot’. The centered map is shown in **Supplementary Figure 2b**. Zero denotes the hotspot location.

**Supplementary Figure 2:**
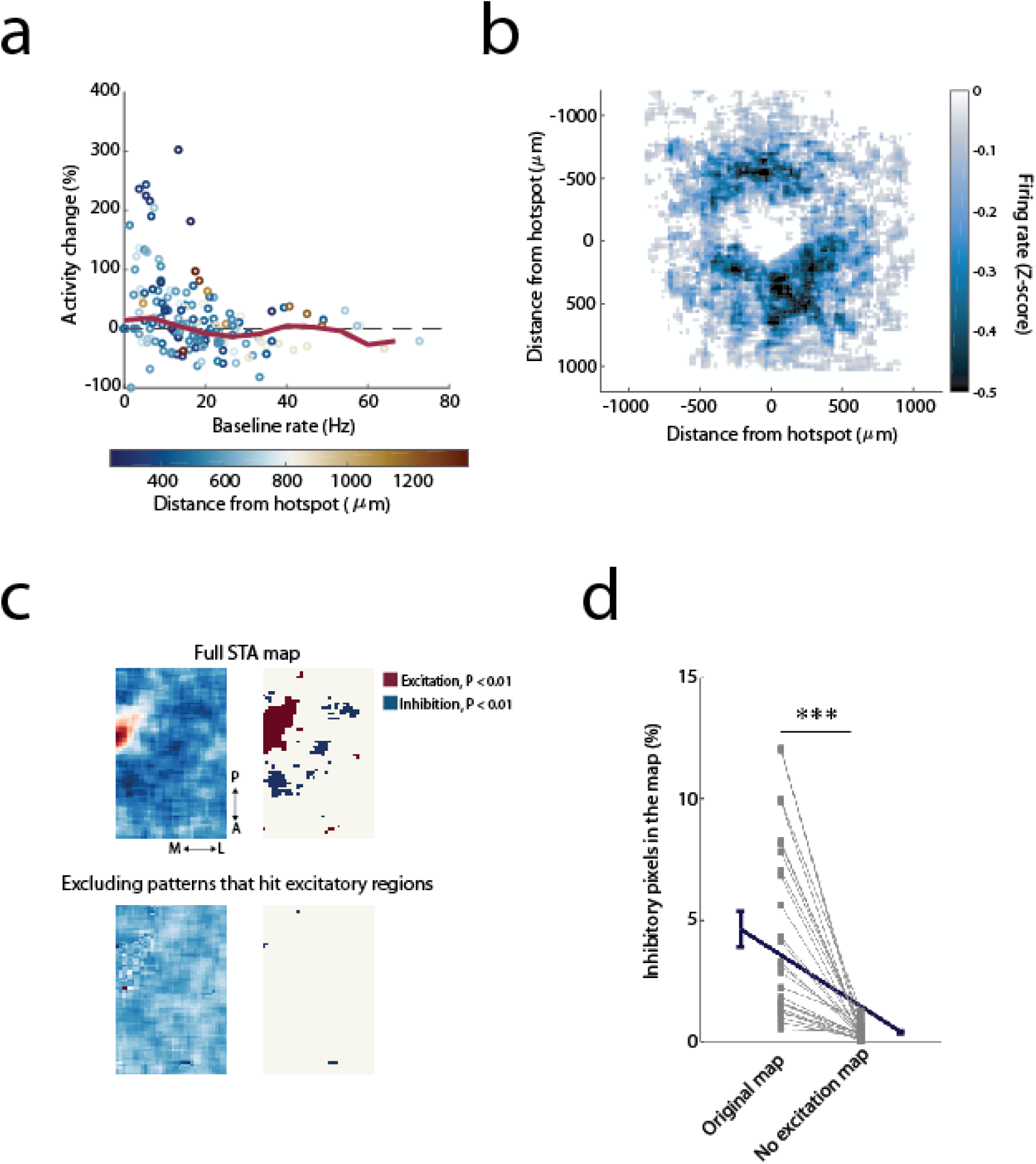
MTC lateral suppression is activity- and spatially-dependent a) Light stimulating a lateral spot without stimulating the hotspot has no effect on the recorded neuron’s baseline firing rate, regardless of its firing rates. Color code as in **Figure 2d**. b) Lateral inhibition is confined in space. All Z-scored STA maps were centered at the hotspot location (N = 27 Z-scored maps, see Methods). c) MTC lateral suppression is effective only when the target MTC is activated. Upper panels: an example of a MTC receptive field map and the corresponding significance map. Lower panels: the same maps recomputed by excluding all light patterns that stimulated the region around the hotspot (All pixels with significant excitation P < 0.01, relative to shuffled data). No inhibitory regions are detected after exclusion. d) Population analysis across all MTC STA maps (N = 27) of the percent of inhibitory pixels in the original STA map and when we excluded the light patterns that hit the hotspot area. A pixel is defined as inhibitory if its value is below two standard deviations from the shuffled distribution (see Methods). The percent of inhibitory pixels drops considerably (P = 1.8e-6, two-tailed paired *t*-test), in the excitation-excluded map.

### Two different neural circuits mediate spikes suppression and entrainment

In our experimental design, the same MTC participated in more than one pairing. Analyzing these pairs revealed that while light stimulation of one pair could enhance the spike precision without affecting its firing rate, activation of a different pair at the same light intensity could suppress the MTC firing rate without affecting its spike precision (Figure 3a). Furthermore, light-activating the same pair with two different intensities could suppress the MTC firing rate at one intensity and enhance spike entrainment in the second (Figure 3b). To quantify these effects at the population level, we compared the firing rate change between pairs that evoked significant inhibition and these that evoked spikes entrainment, and found a significant difference between the two groups (Figure 3c, P < 0.001, two-tailed *t*-test). Consistently, we found only a minor overlap between the groups, suggesting that pairs that exhibit spikes entrainment typically do not exhibit lateral suppression (Figure 3d). These findings suggest that spike suppression and entrainment are not features of the recorded cell type (i.e., mitral versus tufted cells) but instead reflect two different circuits that interconnect MTCs.

**Figure 3:**
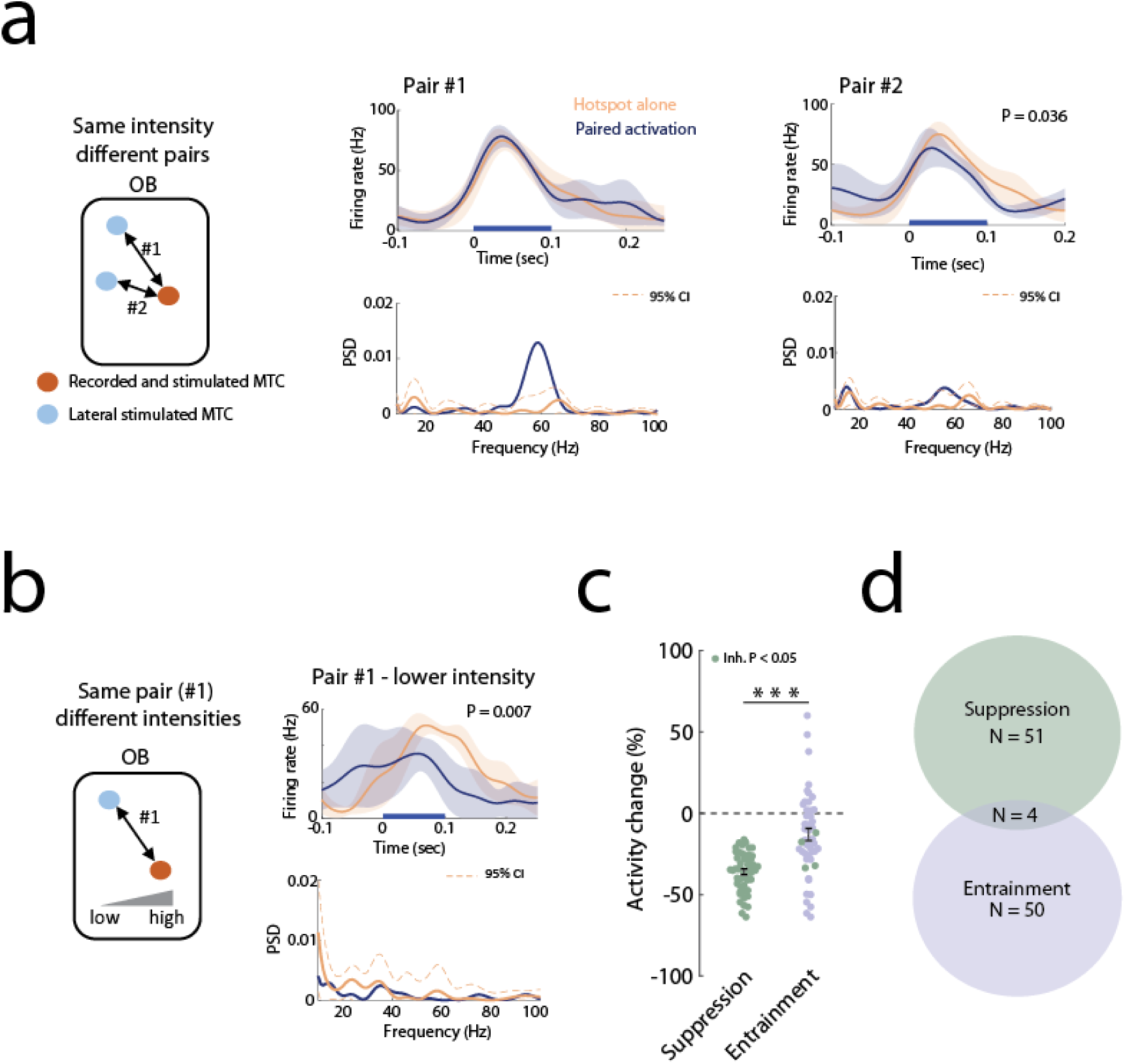
Spike entrainment and suppression are mediated by two different circuits a) Light stimulation of two different MTC pairs sharing the same postsynaptic MTC. Light-activating of pair #1 (left) caused strong entrainment (P < 0.05, two-sample bootstrap) without affecting the light-evoked firing rate (P = 0.59, two-tailed paired *t*-test), whereas light-activating pair #2 (right) suppressed the MTC firing rate (P = 0.036, two-tailed paired *t*-test), without affecting the spikes precision (P > 0.05, two-sample bootstrap). b) Two different light intensities were applied to pair #1, which had differential effects on suppression and entrainment. The high light intensity increased spike entrainment without affecting the firing rate (left panel in **a**). In contrast, lower intensity reduced the light-evoked firing rate (P = 0.007, two-tailed paired *t*-test), with no effect on the spikes’ gamma entrainment (P > 0.05, two-sample bootstrap). c) Mean ± SEM of the activity change following paired activation for pairs that evoked significant suppression (green, N = 51) or spike entrainment (purple, N = 50) in the recorded MTC. Both groups significantly differ in their mean activity change (P < 0.001, two-tailed *t*-test, The shared data points were not included in the statistics to allow two independent samples) Green circles in the entrainment group are pairs that evoked both entrainment and spikes suppression of the recorded MTC firing rates (N = 4). d) A Venn diagram showing a weak overlap between the suppression and entrainment effects.

### Optogenetic activation of GCL neurons increases MTC synchrony in an activity-dependent and location-independent manner

We next sought to understand the neural circuits that mediate spike entrainment and suppression. We examined whether GCL neurons-MTC interactions underlie the temporal changes or the suppressive interactions we found. We conditionally expressed ChR2 in GCL interneurons by injecting AAV5-EF1a-DIO-ChR2 to Gad2-Cre mice into the GC layer (Figure 4a) as in (Dalal and Haddad, 2022; Fukunaga et al., 2014). We used odor stimuli to activate the recorded MTC, while recording MTC spiking activity and the LFP (Supplementary Figure 4a). To optogenetically activate the GCL neurons belonging to a glomerulus column, we used a relatively large light spot sized ∼330 µm^2^ (Egger and Urban, 2006). We then tested how light-activation of subsets of GCL neurons at different locations affects MTC odor-evoked temporal dynamics and spike rate suppression (Figure 4b; N = 4 mice, N = 31 cell-odor pairs from 22 cells). To examine the temporal effects, for each cell-odor pair we extracted the MTC odor-evoked spike phases of the LFP-gamma cycles across all trials. We then quantified the level of spike-LFP gamma coupling using the pairwise phase consistency measure (PPC1). This measure minimizes the bias of the neuron’s firing rate on the spike-LFP synchrony value (Vinck et al., 2012). Only MTCs that were excited by the odor were analyzed (N = 18/31, P < 0.05, two-tailed paired *t*-test). We found that light-activating GC-columns significantly increased MTC spike phase-locking (Figure 4c, N = 3 light spots for each cell-odor pair, a total of 54 light spots tested, P = 0.0016, two-tailed paired *t*-test. Furthermore, within the distance range that we were able to measure, the increased phase-locking did not significantly correlate with the distance from the MTC, and depended on the level of the MTC firing rate (Figure 4d-e). Plotting the odor-evoked spikes of an example high-firing neuron by aligning them to an arbitrary spike in each trial as done in (Fukunaga et al., 2014) further demonstrated that GCL neurons light-activation increased the recorded MTC spikes’ gamma entrainment (Supplementary Figure 4b-c). Our results strongly suggest that MTC-to-MTC lateral entrainment is mediated by spatially distributed GCL neurons.

**Figure 4:**
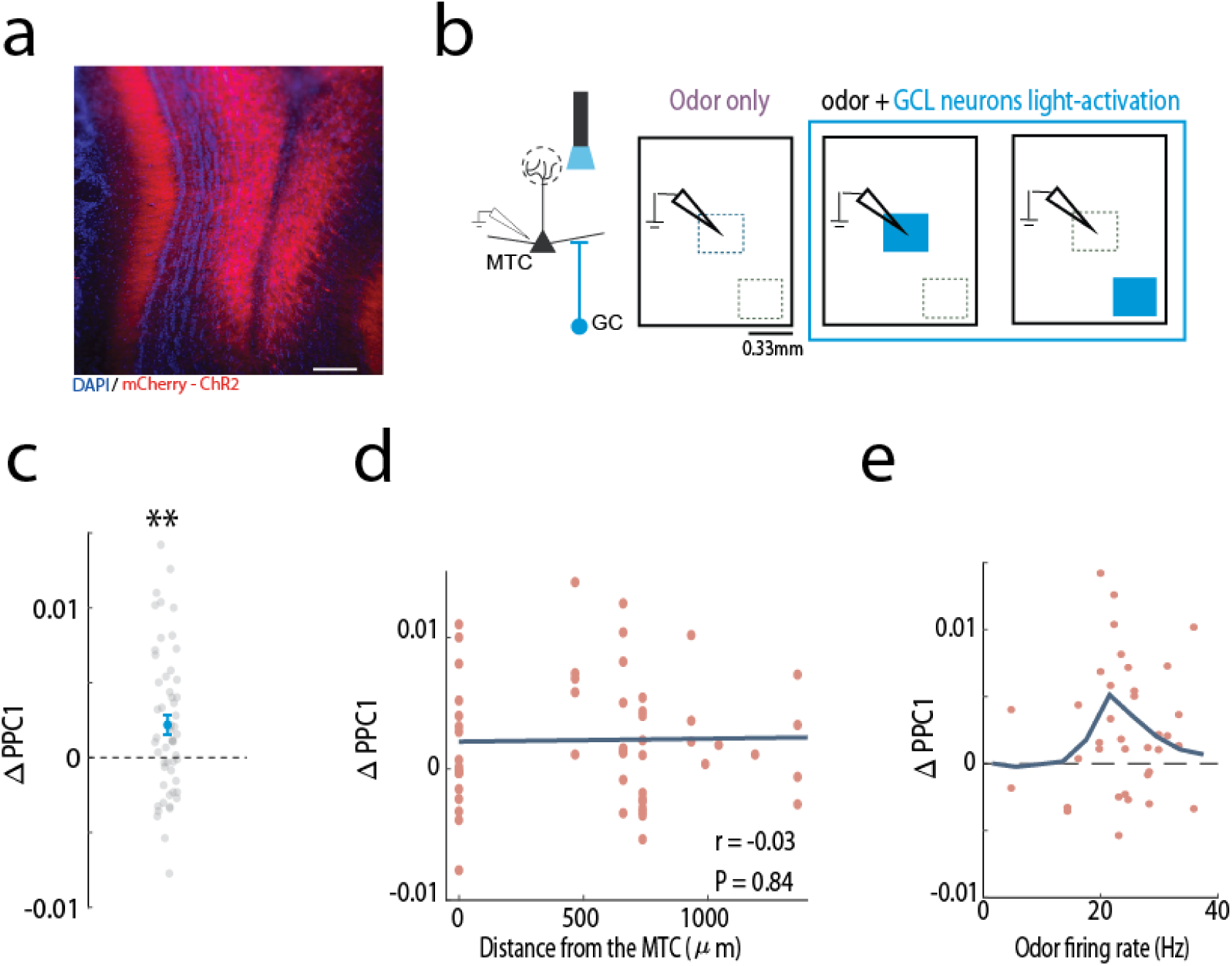
Optogenetic activation of GCL neurons increases MTC synchrony in an activity-dependent and location-independent manner a) Cre-dependent AAV injected into the GC layer (GCL) of Gad2-Cre mice. A coronal OB section showing restricted ChR2 expression in the GCL and the external plexiform layer, into which GCL neurons extend dendrites (red, mCherry-ChR2; blue, DAPI). Scale bar, 0.1 mm. b) Schematic illustrations of the experimental setup. Left: Three weeks post injection, MTCs were recorded while light-activating subsets of GCL neurons. Right: MTC activity was recorded in response to odor stimulation alone (purple) or combined with light-activation of GCL neurons near the recording electrode or distant from it (blue). Scale bar, 0.33mm. c) The change in MTC spike synchrony to the gamma oscillation (L1PPC1) significantly increases under GCL neurons column light-activation during odor stimulation (N = 54, P = 0.0016, two-tailed paired *t*-test). Only cell-odor pairs that were significantly odor-excited were analyzed (N = 18/31 cell-odor pairs; three spots were stimulated per cell odor-pair). d) MTC spike entrainment does not depend on the location of the light-activated GCL neurons. The relation between the change in PPC1 caused by odor and GCL odor stimulation as a function of the distance of the light-stimulated spot from the recording electrode. No significant correlation was found (N = 54 values from 18 cell-odor pairs, r = -0.03, P = 0.84, Spearman correlation). Zero denotes the spot above the recording electrode. e) MTC spike entrainment is activity-dependent. The change in synchrony peaked when MTCs fired at ∼25Hz.

**Supplementary Figure 4:**
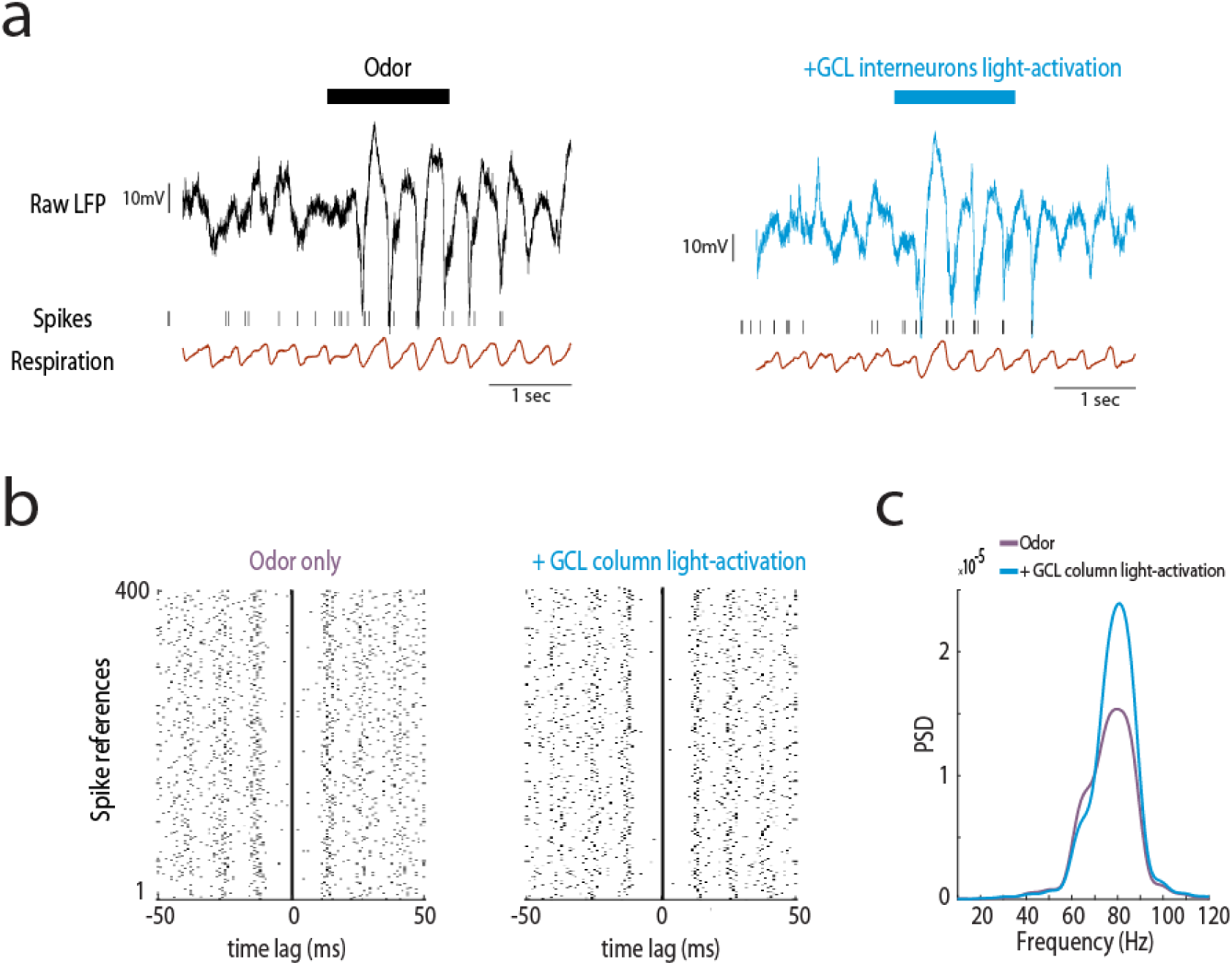
Optogenetic activation of GCL neurons increases MTC synchrony in an activity-dependent and location-independent manner a) Example trials showing the raw LFP data, spike times and respiration signal recorded in response to odor stimulation with and without optogenetic activation of GCL neurons. b) Odor-evoked spike reference analysis. Two spike raster plots are shown, for odor only (left, purple), and odor with light-activation of a GCL neurons column (right, blue). In each raster plot, spikes are plotted relative to a randomly chosen spike during the odor presentation period (N = 400 spikes references, see Methods). Note the potent spike entrainment when GCL neurons are activated. This analysis was performed on a cell that had a sufficiently high firing rate (see Methods). This cell is likely a tufted cell due to its potent entrainment at the high gamma range, as shown in (Burton and Urban, 2021; Fukunaga et al., 2014). c) The power spectral densities (PSD) for the two conditions in **b**. A multi-taper analysis of the circular convolution of each spike raster plot was used to compute the PSD (see Methods).

### MTC suppression is not mediated by GCL neurons

Finally, we examined how GCL neurons optogenetic activation affects MTC odor-evoked firing rate using the same data from the previous experiment (Figure 4). We divided the OB surface into a grid and randomly optogenetically activated GCL neurons columns, with one light spot in each trial of size 330um^2^ (Figure 5a; N = 31 MTCs). We found that MTC baseline activity was strongly suppressed by light activating GCL neurons, mostly when activating neurons in its vicinity (within its column, Figure 5a-b). This experiment validates that in our experimental setup, light stimulation of GCL neurons can suppress MTCs at baseline conditions. In contrast, we found that light-activating sets of GCL neurons during odor stimulation did not change the mean MTC odor-evoked firing rates, irrespective of the MTC firing rate levels. (Figure 5c; N = 54 light spots tested from 18 cell-odor pairs, P = 0.64, two-tailed paired *t*-test). To gain insight into whether the change in MTC odor-evoked response depends on the location of the optogenetically activated GCL neurons, we plotted the activity change as a function of the distance from the recorded MTC (N = 54 light spots tested from 18 cell-odor pairs). There was no relationship between the level of suppression and the location of the activated GCL neurons (Figure 5d). These findings further suggest that GCL neurons are unlikely to underlie the activity-dependent suppression we observed between MTCs (Figure 2). To conclude, our results suggest that MTC-to-MTC lateral entrainment is mediated by spatially distributed GCL neurons, while activity-dependent lateral-suppression, occurs most strongly between adjacent MTCs.

**Figure 5:**
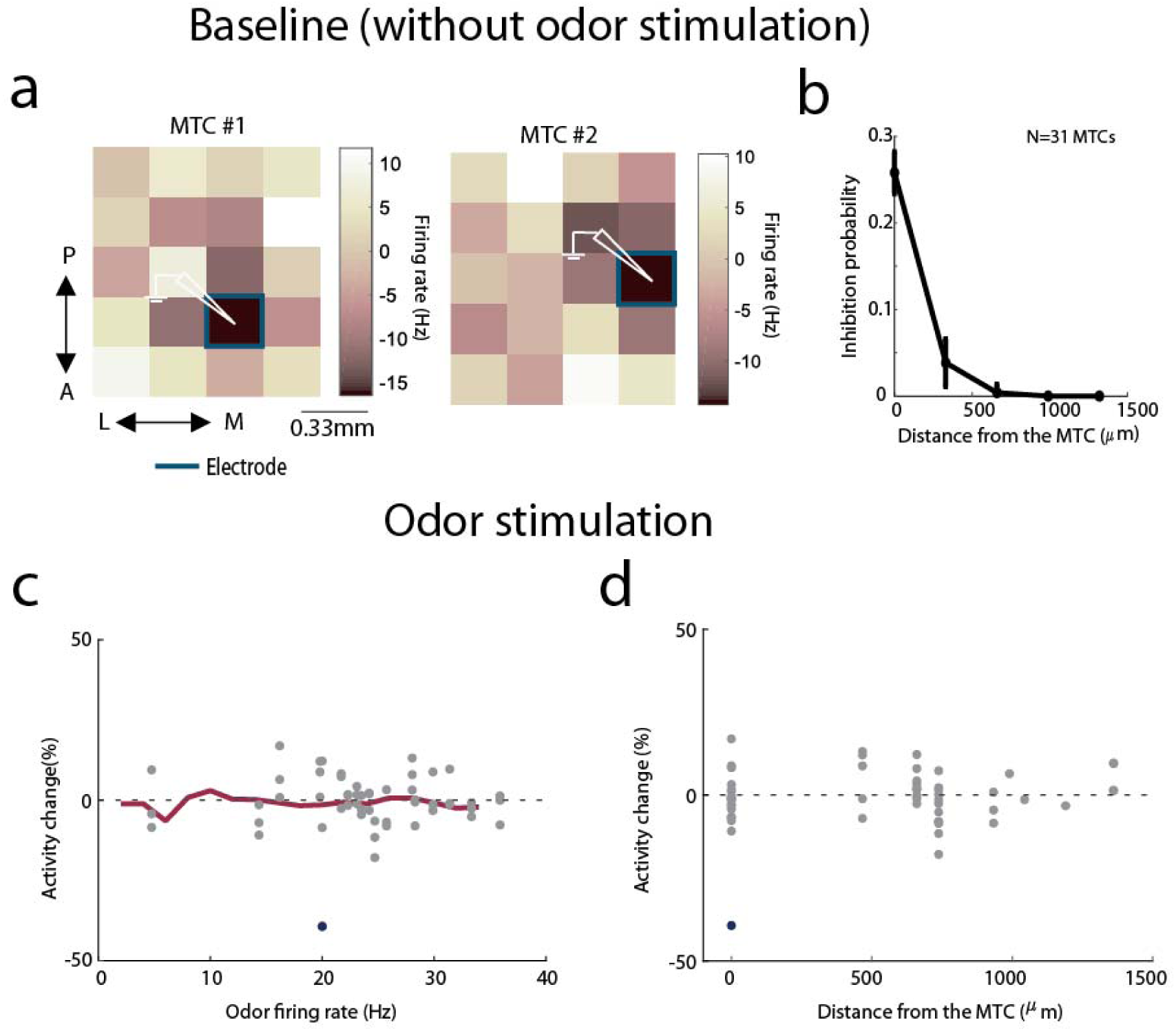
MTC-to-MTC firing rate suppression is not mediated by GCL neurons a) Shown are OB activity maps from two example MTCs, recorded in different mice, during optogenetic activation of GC-columns at baseline conditions. We divided the OB into a grid and activated light spots of size 330 µm^2^. Each entry in the map is color-coded according to the average evoked firing rate across trials. The electrode location is marked by a blue line and a white electrode illustration. b) A population analysis across all OB activity maps (N = 31 MTCs from 4 mice), showing the probability of obtaining a significant inhibition as a function of the distance from the electrode. MTCs inhibition was restricted to activation of GC-columns in their vicinity. c) Similar to Figure 2d, the change in odor-evoked activity following GC activation is plotted as a function of the recorded MTC odor-evoked firing rates (N = 54 values from 18 cell-odor pairs). Filled blue circles denote significant activity change (P < 0.05, two-tailed unpaired *t*-test). A moving average is shown in red. d) Same data shown as a function of the distance from the electrode. Odor-evoked firing rates are not suppressed when a GC column is activated, irrespective of the GC-optogenetic activation location. Zero marks activation of GCs at the location of electrode.

## Discussion

Our findings demonstrate two types of interactions between MTCs that shape their firing rate and temporal dynamics. These two interactions are spatially dissociated: Lateral entrainment spans the whole accessible bulb surface, while lateral suppression occurs primarily between adjacent MTCs. Interestingly, both interactions are activity-dependent, with lateral entrainment and lateral suppression peaking when the postsynaptic MTC fires at ∼40Hz and ∼70-80Hz, respectively. Furthermore, we found that GCL neurons do not mediate substantial MTC-MTC lateral suppression.

### MTC and GCL neurons interactions can facilitate the synchronization of co-active and spatially-dispersed MTCs

It has been shown that two active MTCs can synchronize their stimulus-evoked and odor-evoked spike timings (Doucette et al., 2011; Kashiwadani et al., 1999; Schoppa, 2006). However, how this is achieved and how only the odor-activated MTCs are synchronized is unknown. It has been recently hypothesized that GCs may support the synchronization of odor-responding MTCs via activity-dependent mechanism for GABA release (Egger and Kuner, 2021; Lage-Rupprecht et al., 2020). This hypothesis postulates that the spikes of all strongly co-activated MTCs are synchronized to the gamma rhythm, presumably through the GC network. Consistent with this hypothesis, here we have shown that activation of one group of MTCs can increase the spikes entrainment of another, distant, active MTCs, up to one millimeter away. This finding is consistent with a previous study showing distance-independent synchronization between MTCs (Doucette et al., 2011). We found that this increase in synchronization occurred only if the postsynaptic MTC was firing at a certain level (Figure 1), suggesting a simple mechanism by which only strongly active MTCs are synchronized.

### Lateral suppression is restricted to adjacent and active MTCs

In contrast to the lateral entrainment interactions, we found that lateral suppression of MTC spiking is effective mostly between adjacent MTCs and when the postsynaptic neuron is active at a specific range (∼30-80Hz, Figure 2). Such lateral suppression properties were theoretically suggested (Cleland and Borthakur, 2020; McIntyre and Cleland, 2016; McTavis et al., 2012) and experimentally validated (Peace et al., 2024). We found that in only ∼20% of the tested MTC pairs exhibited significant lateral suppression. This rate is consistent with previous *in-vitro* studies that found lateral suppression between 10-20% of heterotypic MTC pairs (Isaacson and Strowbridge, 1998; Urban and Sakmann, 2002), and is higher compared to a case where the recorded MTC is not active (Lehmann et al., 2016). The role of this lateral suppression, which is most effective when the neurons fire at the gamma range, is unclear. One study suggested that activity-dependent lateral suppression can sharpen MTC odor responses by contrast enhancement (Arevian et al., 2008). However, direct experimental evidence is still lacking.

### GCL neurons do not substantially affect MTC odor-evoked firing rates

GCs are the most abundant interneuron type in the OB (Shepherd, 1972), suggesting they have a key role in regulating MTC activity. In line with this hypothesis, several key studies have provided evidence that GCs may suppress the MTC firing rate (Arevian et al., 2008; Giridhar et al., 2011; Yokoi et al., 1995), while later studies demonstrated that EPL interneurons have a key role in suppressing MTC activity (Burton et al., 2024; Huang et al., 2013; Kato et al., 2013). Here, we show that optogenetically activating GC columns during odor stimulation did not substantially affect MTC odor-evoked firing rates. Although this contradicts a previous in-vitro result (Arevian et al., 2008), it is consistent with a recent in-vivo study that found no increase in MTC firing rates when silencing GC activity either in anesthetized or in awake mice (Fukunaga et al., 2014). It is worth noting that light-activating large numbers of GCs can suppress MTC odor-evoked responses (Dalal and Haddad, 2022; Gschwend et al., 2015). However, activation of all GCL neurons is unlikely to occur during natural odor response. We speculate that MTC-to-MTC suppression is mediated by EPL neurons, most likely the Parvalbumin neuron (PV). This hypothesis is based on their activity and connectivity properties with MTCs (Burton, 2017; Burton et al., 2024b; Guest et al., 2021; Huang et al., 2013; Kato et al., 2013; Miyamichi et al., 2013). Of note, it is possible that our viral transfection protocol in Gad2-Cre mice might not transfected all subtypes of GCs, which may explain the lack of substantial suppressive effect onto MTCs. Future studies are required to shed light on how PV neurons affect MTC activity.

### MTC-GCL neurons interactions under anesthesia

Our experiments were performed under Ketamine anesthesia, an NMDA receptor antagonist that affects the reciprocal dendro-dendritic synapses between MTCs and GCs (Egger and Kuner, 2021; Lage-Rupprecht et al., 2020). Consistent with that, recent studies reported lower GC excitability under anesthesia (Cazakoff et al., 2014; Kato et al., 2012). This raises the concern that our result might not be valid in the awake state. We argue that this is unlikely. First, (Fukunaga et al., 2014) reported that GCs baseline activity in anesthetized and awake mice is similar, suggesting that MTC-GC synapses function. Second, we show that light activation of GCL neurons strongly inhibits the MTC baseline activity (Figure 5) and increases MTC odor-evoked spike-LFP coupling in the gamma range (Figure 4). These experiments validate that GCL neurons can exert inhibition over MTCs in our experimental setup. Third, we have shown that light-activating all accessible GCL neurons has a minor effect on the MTC odor-evoked firing rates in an awake state (Dalal and Haddad, 2022), corroborating the finding that GCL neurons are unlikely to provide strong suppression to MTCs. Fourth, and most importantly, we showed that optogenetic stimulation of MTCs entrains other MTC spike times, which is achieved via the GCL neurons. This suggests that the lack of lateral suppression following MTC or GCL neuron opto-activation is not due to MTC-GC synapse blockage. That said, we cannot exclude the unlikely possibility that NMDA receptor blockage under anesthesia impairs MTC-to-MTC suppressive interactions but not the MTC-to-MTC mediated spike entrainment.

### Downstream integration of synchronous MTC activity

How do downstream olfactory cortex neurons benefit from synchronous MTC activity? Anterior piriform cortex (aPC) neurons integrate the activity of several active MTCs and act as coincidence detectors (Davison and Ehlers, 2011; Haddad et al., 2013). Recently, we have shown that increased MTC spike phase-locking to OB gamma oscillations enhanced aPC neurons’ odor representation (Dalal and Haddad, 2022). This current study extends these findings as it demonstrates that the co-activation of MTCs increases their spikes’ gamma entrainment. Moreover, this lateral synchronization is mediated via the GC network. In summary, we found that MTC interactions shape odor-evoked MTC firing rates and spike times. Both of these changes are activity-dependent. Activity-dependent synchronization can enable the synchronization of odor-activated MTCs dispersed across the glomerular map. Since glomeruli columns are zero-phase synchronized (Peace et al., 2024), and the odor-activated MTCs are synchronized to the same phase (Doucette et al., 2011; Kashiwadani et al., 1999), the efficacy of odor information transform to downstream neurons will increase (Dalal and Haddad, 2022). Moreover, activity-dependent spike suppression might enhance odor representation in the OB by reducing low and noisy odor-evoked MTC responses. Finally, our findings suggest that two types of OB interneurons mediate these distinct interactions.

## Acknowledgments

This study was supported by a grant from the Israel Science Foundation ICORE program [11/51].

## Author Contributions

T.D. performed the experiments and analyzed the data, T.D. and R.H conceptualized the experiments and wrote the paper.

## Methods

All surgical and experimental procedures were conducted in accordance with the National Institutes of Health Guide for the Care and Use of Laboratory Animals and Bar Ilan University guidelines for the use and care of laboratory animals in research, and were approved and supervised by the Institutional Animal Care and Use Committee (IACUC). Animals were housed in a group cage and received no experimental treatment, except genotyping. The animals were maintained in a reverse light/dark cycle, and all experiments were performed during the dark period. 19 Tbet-Cre (Jackson Laboratory, Stock No. 024507) crossed with Ai32 (Jackson Laboratory, Stock No. 012569), and 4 Gad2-ires-Cre (Jackson Laboratory, Stock No. 028867) male and female mice aged 3-12 months were used. As no differences in light-evoked responses were observed between sexes, data from both sexes was pooled.

### Surgical Procedures for Electrophysiology Recordings

Animals were anesthetized with ketamine/ medetomidine (60/ 0.5 mg/ kg, i.p.) and then fixed in a stereotaxic frame. The bone overlaying the dorsal OBs was removed. Additional anesthesia was administered as needed (∼30% of the original dose of ketamine/ medetomidine). Body temperature was maintained at 36-37°C using a homoeothermic blanket system (Harvard Apparatus).

### Viral Injections

Mice were briefly anesthetized with isoflurane before an intraperitoneal injection of ketamine/ medetomidine (60/ 0.5 mg/ kg i.p.). The mouse was head-fixed in a stereotaxic frame, and AAV5-EF1a-DIO-ChR2-eYFP/mCherry (titers: 4×10^12^ particles/ ml; University of North Carolina Gene Therapy Center) was injected into the granule cell layer. To infect GCL neurons, we modified an injection protocol employing five injection sites in two tracks, as reported elsewhere (Fukunaga et al., 2014). The coordinates for the first track were +0.9 mm (M-L) from the midline rhinal fissure, +0.8 mm (A-P), and three sites at the D-V axis at -0.8 mm (300 nl), -1.1 mm (200 nl), and -1.3 mm (200 nl). Second track coordinates were +0.9 mm (M-L) from the midline rhinal fissure, +1.2 mm (A-P) and 2 sites at the D-V axis at -0.8 mm (200 nl), and -1.1 mm (200 nl). M-L coordinates are relative to the midline rhinal fissure. The syringe was left in place for at least three minutes before moving to the next coordinate on the D-V axis. The virus was injected using a micro-injector (IMS-10, Narishige, Japan) at a rate of 70 nl/ min, which was left in place for 5 minutes to allow viral particle diffusion before needle removal. Incisions were closed with tissue glue (Vetbond), and an analgesic injection (Carprofen) was administered at the end of the surgery. Electrophysiology was carried out at least three weeks post-viral injection.

### Electrophysiology

The neurons’ extracellular activity and local-field potential (LFP) were recorded using tungsten electrodes (∼1-10 MΩ; FHC). Neural signals were amplified and first filtered at 1-10,000Hz and then at 300-5,000 Hz for spiking activity (AM-Systems 1800), sampled, and recorded at 40 kHz (National Instruments, Austin, TX). Spike signals were sorted offline using MClust software in MATLAB (written by A.D. Redish, University of Minnesota). MTC recordings were collected from the dorsal OB (∼200-500 µm). The electrode was typically lowered at an angle of 90 degrees. The depth of recorded neurons was estimated when the recording electrode was withdrawn using a micromanipulator.

### Optogenetic Stimulation of the Olfactory Bulb

Optogenetic stimulation of neurons in the OB was performed using an optical imaging system based on a digital micro-mirror (OPTOMA X600 DLP Projector), as described in (Grobman et al., 2018). Precise spatial control of optical stimulation was achieved by projecting two-dimensional light patterns over the dorsal surface of the OB. We used blue and white light for MTC and GC activation, respectively. For MTC stimulation (Tbet-Cre mice crossed with Ai32 mice), a blue filter was placed on top of the collimating lens, which was placed at a distance of ∼20 cm from the projector (f = 75 mm, Achromatic doublets, Thorlabs). This configuration resulted in an image where each projected pixel corresponded to a square of 22 µm^2^. The size of the craniotomy determined the light stimulation boundaries of each experiment. Optical stimulation was controlled with the MATLAB psychophysical toolbox. We used a photodiode (FDS1010, 400-ns rise time, Thorlabs) to obtain a timestamp for each light stimulus. The light intensity ranged between ∼0.1-∼1.5 mW/ mm^2^ as measured with an optical power meter (Thorlabs PM100D). The stimulation frequency of the optical imaging system was 120 Hz.

### Spike-Triggered Average (STA)

The spike-triggered average calculation was described previously by (Grobman et al., 2018). Briefly, STA was used to characterize the receptive field of the recorded MTC. Here, we use the term ‘receptive field’ in a more abstract sense to refer to the ensemble of all neurons on the bulb’s surface that modify the recorded neuron activity.

We stimulated the dorsal OB with patterns of multiple square spots measuring 88-110 µm^2^. Light spots within a pattern could overlap up to a shift of one pixel (22µm). The stimuli duration was 0.1 sec, and the inter-stimuli-interval between consecutive patterns ranged between 0.1-0.2 sec. The number of spots projected in each trial ranged between 5-10, and the number of trials per recording ranged between 2000-3000. We computed the response map for each cell by computing the spike-triggered average, which is the weighted firing rate average of all projected stimuli.

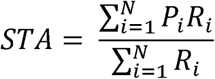

P_i_ is the 2-dimensional projected light pattern, and R_i_ is the evoked firing rate following stimulus P_i_.

### Paired light stimulation

Single light spots (over the hotspot or a lateral spot) or paired stimulation (both together) of size ∼154 µm^2^ were projected at varying light intensities (range ∼0.1 to ∼1.5 mW/ mm^2^, reported in Figures 1-3). Spots were selected either randomly or manually. In the manual selection case, we selected spots that caused either significant or mild but insignificant inhibitory effects on the recorded MTC (e.g., local cold spots in the receptive-field map; see example in Figure 2a of example spots that were selected manually). The number of trials for each condition at a given light intensity was 15, and the inter-stimuli interval (ISI) was set to 1.5 seconds.

### Optogenetic activation of GCL neurons during baseline and odor stimulation

To verify the effect of optogenetic activation of GCL neurons on the recorded MTC, we scanned the OB with light spots, without stimulating the MTC. Light spots were of size 330µm^2^, each spot was illuminated 20-30 times, the light duration was 0.2 sec, and the ISI was 0.4 sec. The light intensity was as described in (Dalal & Haddad 2022). As expected, we found a ‘coldspot’ (i.e. a reduction in firing rate) in the vicinity of the electrode location (Figure 5), confirming that light-activating GCL neurons evokes robust inhibition to reduce MTC baseline activity, consistent with a recent study (Huang et al., 2016). For optogenetic activation of GCL neurons during odor stimulation, the odor duration was 1.5 seconds. The light stimulus was active throughout the odor stimulation period. The activation of GCL neurons at different locations during odor stimulation was performed based on the activity map generated under baseline conditions. In each of the three conditions in this experiment, odor stimulation alone or combined with GCL neurons column activation, the number of trials was at least 10, with an ISI of 10-15 seconds. In these experiments, the LFP was simultaneously recorded using the same electrode, and the signal was later filtered for spiking activity (300-5000 Hz) and LFP ranges (1-300 Hz).

### Respiration Analysis

We recorded the respiration signal using a piezoelectric sensor (APS4812B-LW100-R, PUI Audio). The most salient feature of the respiratory signal is the peak in the middle of the inhalation cycle caused by the pressure of the diaphragm on the sensor. The onsets of inhalation and exhalation were defined as the zero-crossings of the signal before and after the peak, respectively.

### Odorant Application

Odorants were applied using a custom-built olfactometer. Odorants were diluted in mineral oil (1:100) and stored in sealed glass vials. This concentration was chosen to elicit a detectable response. The tubes were placed in front of the animals’ nostrils at a distance of ∼2 cm. Airflow was controlled with a mass flow controller (Agilent, Alimc-2LSPM) and set to 0.8 slpm. Air circulated freely between stimulations to reduce odorant remnants. A vent removed residual odorants. Odorant stimulation times and sequences were controlled by custom MATLAB scripts. The odorant stimulation time was set to 1.5 sec., with an inter-trial interval of 10-15 sec. The odorant sequence was randomized, and each stimulus was delivered at least ten times. All odorants used (all at 1% dilution) were from Sigma-Aldrich at their highest purity. The odorants used were Phenethylamine (PEA (CAS: 64-41-0), Ethyl acetate (CAS: 141-78-6), Ethyl valerate (CAS: 539-82-2), 2-heptanone (CAS: 110-43-0), Ethyl butyrate (CAS: 105-54-4), Ethyl tiglate (CAS: 5837-78-5), and Acetophenone (CAS: 98-86-2).

## QUANTIFICATION AND STATISTICAL ANALYSIS

### General Statistical Analysis

All data analyses were performed in MATLAB. The figures or figure legends detail the number of data points used for all statistical tests and graphs. Significance alpha was defined as a 0.05/0.01. Unless stated otherwise, we report the mean ± SEM or %95 confidence interval. Mean ± standard deviation is reported when estimated from a bootstrap process. We used a *t-*test as required by the test’s null hypothesis and population assumptions. All tests were two-tailed. The test is reported in the figure legend or the main text.

### Data Analysis Spike Sorting

Spike signals were sorted and clustered offline using MClust software in MATLAB (written by A.D. Redish) or Spike3D (Neuralynx). Only visually well-isolated clusters were used, with less than 5% of spikes violating an inter-spike interval of 2 ms.

### Significant regions in the STA map

Computation of significant activity change of each pixel of the STA map was done relative to a shuffled STA map. The shuffled map was generated by shuffling the patterns indices and computing the shuffled STA (multiplying the shuffled patterns with the original firing rate as described above). Then, the 0.5 and 99.5 percentile values from the shuffled map were determined. The values in the STA map that were above or below these high and low percentile thresholds were assigned as significant excitatory and inhibitory pixels, respectively.

### Superimposed STA maps

To analyze the spatial location of the inhibitory regions surrounding the recorded MTC, each STA map was Z-scored using the mean and standard deviation values on the map. We then thresholded the map such that only pixels with a Z-score <= -1 remained. The hotspot peak was centered at the origin, and the inhibition values were plotted on the map relative to the hotspot. We superimposed all maps and averaged across all neurons. We computed the radial average of this map by averaging all angles from the center of the map.

### Exclusion of excitatory regions in STA maps

STA maps were recomputed by excluding trials with at least one light spot that hit the significant excitatory regions in the map. The significance of each pixel in the recomputed map was registered, and the percent of inhibitory pixels in the original and recomputed map was assigned. To verify that the lower number of significant pixels in the recomputed map was not due to a smaller number of trials, for each full map we randomized the same number of trials that were used to construct the recomputed map and computed the STA. We found no significant change in the percent of inhibitory spots compared with the original map (data not shown).

### Quantification of MTC lateral inhibition

A pair of light-stimulated MTCs was considered as evoking lateral inhibition if it met two criteria: the response to the hotspot stimulation was significantly different from baseline (P < 0.05, one-tailed paired *t*-test), and there was a significant change in the firing rate across trials between hotspot stimulation and paired activation within a window of 200 ms (P < 0.05, two-tailed unpaired *t*-test, stimulus duration 100 ms). A window of 200 ms was used to account for neural responses with a slower return to baseline. The distance between the spots in the pair was measured using Euclidean distance. The activity change measure was defined as:

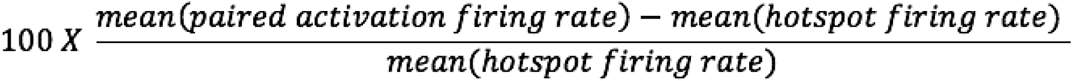

### MTC entrainment across trials

Quantifying MTC spike entrainment across trials was done by computing the PSTH of each condition during the light period (100 ms) using a Gaussian kernel with a standard deviation of 2 ms. The PSTHs were zero-padded to a length of 1 second, Z-scored, and the power spectrum density was computed using multi-taper analysis (TW = 3 Hz, L = (2*TW)-1, where L is the number of orthogonal Slepian tapers). The change in entrainment measure was the difference between the integral of paired activation and the hotspot alone at 40-70 Hz. We analyzed only pairs and light intensities that significantly responded to light stimulation (P < 0.05, two-tailed paired *t*-test). Shuffled distribution of spike entrainment was obtained by shuffling the trial identities and computing the power spectrum density integral difference. The differential entrainment values were labeled significant if they crossed the 95% confidence interval around the shuffled distribution.

### Time-Frequency Analyses of MTC entrainment

Time-frequency representation was performed using the continuous wavelet transform (the analytic ‘Morse’ wavelet, 8 octaves, 32 voices per octave; MathWorks) over the Peri-stimulus time histograms (PSTH) across trials (Gaussian window of 2 ms standard deviation).

### Odor-Evoked Responses

An odor firing rate was computed over the first two seconds following odor presentation. This time window included an additional 0.5 sec after stimulus offset to account for neural responses with a slower return to baseline. Trials were aligned to odor onset so that the analyzed neural activity would be aligned with the light stimulation. A response was defined as significant if the firing rate was significantly different from the average firing rate over an equivalent time window prior to stimulus onset (P < 0.05, two-tailed paired *t*-test). PSTHs were smoothed using a Gaussian filter (standard deviation of 50 ms, Chronux toolbox).

### LFP Analysis

Recorded LFP signals were down-sampled to 4 kHz and band-pass filtered to the γ-band frequency range (40-70 Hz) using the MATLAB (MathWorks) filter designer (Butterworth IIR filter with order 1). The line noise (50 Hz) was filtered using an IIR bandstop filter (48-52 Hz). We focused our analysis on the low γ-band frequency range as a prominent oscillation in this range in the olfactory bulb was previously reported (Cenier et al., 2009; Lepousez and Lledo, 2013).

### Spikes-LFP pairwise phase consistency (PPC)

Spike-LFP coupling was computed using the pairwise phase consistency 1 (PPC1) measure (Vinck et al., 2012, 2013). For each neuron, we extracted the spike phases from all trials. To compute the spike phases per trial, we extracted the instantaneous phase of the corresponding γ-band-filtered LFP (40-70 Hz) using the Hilbert transform, assigned a phase to each spike (0–2π) and pooled the phases across all trials. The PPC1 values were the mean dot-product of all pairwise spike phases, excluding phases of spikes from the same trial.

### Spike reference analysis

We performed a spike reference analysis to quantify the entrainment of odor-evoked MTC spikes, as seen elsewhere (Fukunaga et al., 2014). Spike reference raster plots were constructed for each condition by randomizing 400 spikes from all trials that occurred at a time window of 1 second after odor onset. Each randomized spike served as a reference and was set to time lag 0, and the spikes in a window of ± 50 ms around it were plotted relative to the reference. Choosing a different number of randomized spikes or a different window size did not affect the results. We then computed the PSTH of the raster plot, subtracted its mean, and performed a circular convolution. To compute the power spectral density of the raster plot, we zero-padded the convolved signal into one 1-second length and applied a multi-taper analysis (TW = 5 Hz). The PSD curve was smoothed using a rectangular window of order 15.

